# Intra-Host Mutation Rate of Acute SARS-CoV-2 Infection During the Initial Pandemic Wave

**DOI:** 10.1101/2023.03.24.534062

**Authors:** Kim El-Haddad, Thamali M Adhikari, Tu Zheng Jin, Yu-Wei Cheng, Xiaoyi Leng, Xiangyi Zhang, Daniel Rhoads, Jennifer S. Ko, Sarah Worley, Jing Li, Brian P. Rubin, Frank P. Esper

**Author notes:** **<u>Corresponding author:</u>** Kim El Haddad, MD, Address: R3, 9500 Euclid Avenue, Cleveland, Ohio 44195 USA. **<u>Alternative Corresponding author:</u>** Frank Esper, MD, Address: R3, 9500 Euclid Avenue, Cleveland, Ohio 44195 USA.

## Abstract

**Background:** Our understanding of SARS-CoV-2 evolution and mutation rate is limited. The rate of SARS-CoV-2 evolution is minimized through a proofreading function encoded by *NSP-14* and may be affected by patient comorbidity. Current understanding of SARS-CoV-2 mutational rate is through population based analysis while intra-host mutation rate remains poorly studied.

**Methods:** Viral genome analysis was performed between paired samples and mutations quantified at allele frequencies (AF) ≥0.25, ≥0.5 and ≥0.75. Mutation rate was determined employing F81 and JC69 evolution models and compared between isolates with (ΔNSP-14) and without (wtNSP-14) non-synonymous mutations in NSP-14 and by patient comorbidity.

**Results:** Forty paired samples with median interval of 13 days [IQR 8.5-20] were analyzed. The estimated mutation rate by F81 modeling was 93.6 (95%CI:90.8-96.4], 40.7 (95%CI:38.9-42.6) and 34.7 (95%CI:33.0-36.4) substitutions/genome/year at AF ≥0.25, ≥0.5, ≥0.75 respectively. Mutation rate in ΔNSP-14 were significantly elevated at AF>0.25 vs wtNSP-14. Patients with immune comorbidities had higher mutation rate at all allele frequencies.

**Discussion:** Intra-host SARS-CoV-2 mutation rates are substantially higher than those reported through population analysis. Virus strains with altered NSP-14 have accelerated mutation rate at low AF. Immunosuppressed patients have elevated mutation rate at all AF. Understanding intra-host virus evolution will aid in current and future pandemic modeling.

## Background

Since the introduction of the SARS-CoV-2 pandemic in 2020, over 102 million cases have been reported within the United States (1). During this time, multiple variants have emerged associated with alteration in clinical outcomes, disease severity and transmission dynamics (2). SARS-CoV-2 rate of mutation are commonly estimated through inferring substitution rate matrix based on phylogenetic tree using maximum likelihood methods through analysis of global databases comprised of unrelated virus sequences submitted ad hoc(3,4). This population-based rate began at a modest 21.9 substitutions/genome/year in the initial months but has steadily risen over the course of the pandemic where it is now estimated at ∼28.4 substitutions/genome/year (5). However, viral mutation rate during the course of the infection remains poorly understood with few studies describing intra-host kinetics.

Analysis of SARS-CoV-2 mutations within a host during the course of an infection have been highly variable and are affected by sequencing protocols and data analysis parameters(i.e. variant-calling) (6,7). The mutation rate of SARS-CoV-2 genome is slower than most RNA viruses predominantly through the action of nonstructural protein 14 (NSP-14) (8). NSP-14 is present in all coronaviruses and contains an *N*-terminal ExoN domain providing replication fidelity for the RNA dependent RNA polymerase important for viral replication and transcription (9–11). Mutagenesis of NSP-14 enzymatic activity is thought to have significant impact on increased genomic mutation diversity (12). ExoN inactivation was shown to create a “mutator phenotype,” leading to a 15- to 21-fold rise in mutations during replication in cell culture but may adversely affect viral fitness (10).Additionally, viral mutagenesis is reported to be influenced by host comorbidities (13). Subsequently, there is concern that novel variants eliciting immune escape emerge within immunocompromised hosts following prolonged infection (7).

To better understand the mutation capacity of SARS-CoV-2, we perform analysis of paired samples and calculate the intra-host mutation rate with further examination of the effects of altered NSP-14 and host comorbidity. Better insight on this viruses ability to evolve has importance for both current and future coronavirus pandemics (14).

## Methods

### Sample Identification and collection

Patient samples were identified through The Cleveland Clinic Pathology and Laboratory Medicine Institute (PLMI) SARS-CoV-2 variant surveillance project(2). Selected samples focused on the period of the initial pandemic wave between 3/17/2020 and 5/27/2020. This period was chosen as treatment was limited and immune-preventative strategies (e.g. immunizations, monoclonal antibodies) against SARS-CoV-2 were not available. Additionally, SARS-CoV-2 re-infection was unlikely during this period. Hence, the mutation rate analysis is unlikely to be influenced by these external factors.

Adults age ≥ 18 years with multiple positive nasopharyngeal samples occurring within 5 to 60 days of initial screening were identified. This interval time frame was selected to prevent skewing of model results from short sampling intervals while further minimizing chance of re-infection with different SARS-CoV-2 strains (15,16). Only pairings where initial and subsequent samples had cycle threshold (CT) ≤ 30 were included to ensure high quality genomic sequencing. Children <18 years were excluded as identification of SARS-CoV-2 in children during the first wave was minimal. Those specimens with an indeterminate result, obtained from locations other than the nasopharynx, or whose samples contained discordant viral lineages (suggesting reinfection) were also excluded.

Patient comorbidities were identified through the COVID-19 registry (17). Patients were classified into four comorbidity categories: Endocrine (obesity and diabetes mellitus), cardiac (hypertension and coronary artery disease), pulmonary (asthma, obstructive sleep apnea and COPD) and immunologic (autoimmune diseases, history of prior/current cancer and current immunosuppression therapy). Sample collection and medical review is approved by the Internal Review Board at Cleveland Clinic.

### Library preparation and sequence data analysis

Following patient identification, initial and subsequent nasopharyngeal samples were retrieved from Biobank freezers housed at PLMI and processed for viral genome analysis though next generation sequencing (NGS). Total nucleic acids were purified from each specimen and subjected to reverse transcription (RT), NGS library preparation, sequencing, and data analysis according to the manufacturer’s recommendation (Paragon Genomics, Hayward CA). Briefly: Total RNA from SARS-CoV-2 was converted into complementary deoxyribonucleic acid (cDNA) synthesis via RT in 20 μL reactions (10 minutes at 8°C and 80 minutes at 42°C). The derived panel of 343 amplicons utilized for SARS-CoV-2 enrichment covers 99.7% of the viral genome (MN908947/NC_045512.2) with 92 bases uncovered at each end. Purified cDNA was subject to multiplex PCR (10 minutes at 95°C, followed by 10 cycles at 98 °C for 15 seconds each and 60 °C for 5 minutes). Excess primers and oligos were subsequently removed from the purified PCR products, after which a second round of PCR to append indexing primers was performed (initial denaturation, 10 minutes at 95°C, followed by 24 cycles of 98°C for 15 seconds and 60°C for 75 seconds). Sequencing libraries were then prepared and quality was assessed visually using an Agilent® 2100 Bioanalyzer® (Agilent, Santa Clara CA). The presence of a ∼275 bp peak indicated successful amplification and these libraries were then sequenced using a MiSeq instrument (Illumina, San Diego, CA). Raw fastq reads was extracted by Illumina bcl2fastq (v2.20.0) and mapped to the reference genome Wuhan-Hu-1 (NC_045512.2) using BWA program (18). Variants were called using FreeBayes program (19) and filtered at 5% and 10% allele fractions for insertion or deletion (INDEL) and single nucleotide variants (SNV), respectively. Amino acid changes were annotated using snpEff (v4.5) program (20). All variant data was visually examined in Integrative Genome Browser (IGV, version 2.11.0) (21) to eliminate artifacts. Quality was ensured by monitoring mapping quality, phred score, and manual review.

### Variant Calling

Variant calling methodology is strongly dependent on the library protocol and sequencing technology and requires tuning of parameters to distinguish true variants from false positive calls (22). Variant calling was expanded from established WHO criteria (23) and was performed by manual review of each SNV by three independent investigators through IGV (21). We used a minimum depth of ≥100 reads at each position for all samples and quantified SNV at 3 separate allele frequencies (AF ≥0.25, AF ≥0.5, and AF ≥0.75). AF was defined as the proportion of SNV in the sample reads. Mutation change represents the discordance in SNVs between initial and the subsequent samples at each AF. In addition, SNVs below 0.25 AF and those mutations where investigator consensus was not achieved were excluded from the analysis to ensure no overestimation of mutation rate. Following classification of mutation (missense, silent, nonsense, INDEL) and location within the genome, isolates with non-synonymous mutations of NSP-14 were identified and placed in the ΔNSP-14 group. As our understanding of SARS-CoV-2 NSP-14 is evolving, no weight was given to mutation types (Missense vs frameshift vs nonsense) or location within NSP-14 (active vs structural site). Changes in genome between initial and subsequent samples were quantified for each pair and used for calculation of mutation rate (standardized to mutations/genome/year) through both F81 and JC69 models (below).

### Calculation of Genome Mutation rate

We chose two mutation models (F81 and JC69) in calculating the overall substitution rates between samples (24,25) as sample size was limited and both models assume equal mutation rates across different nucleotides allowing for a smaller number of model parameters. JC69 also assumes equal base frequencies, whereas F81 allows for variable base frequencies with equal substitutions providing a more realistic calculation of the mutation rate. For both models, mutation rates were estimated by the use of maximum likelihood algorithms. Hereafter, the results detail findings from the F81 model while results detailing findings from the JC69 analysis appear in the supplementary materials.

### F81 model derivation

For each of the *n* patients, we obtained two virus specimens at different time points and the time interval is denoted as *t*_k_ for patient *k*. To obtain the maximum likelihood estimate of the mutation rate based on the evolutionary model F81, we assume all the patients are independent. Therefore, the likelihood of the data (*L*) is the product of the likelihood (*L*_k_) of each patient *k*, measuring the probability of observing the sequence evolving over time *t*_k_. Because for each patient, both initial and subsequent sequences were available, under the assumption that all the nucleotides are independent, the probability *L*_k_ is the product of the probability over all nucleotides. Under the model F81, the probability that a nucleotide *i* (*i* ∈{A, T, G, C}) remains unchanged over time *t* is

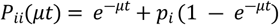

and the probability of a nucleotide *i* to change to a nucleotide *j* over time *t* is

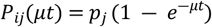

where *u* is the mutation rate per nucleotide per year, and *p*_*i*_ is the frequency of nucleotide *i*. Let *l*_(ij),k_ denote the number of nucleotide *i* changed to nucleotide *j* for patient *k* (in the case of *i* is the same as *j*, the nucleotide remains unchanged), the overall likelihood can thus be represented as

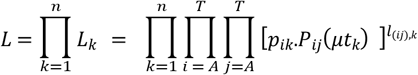

where *p*_*ik*_ is the frequency of nucleotide *i* in the first specimen of the *k*^th^ patient (in practice, these frequencies are very similar to the frequencies from the SARS-CoV2 reference sequence). The log likelihood is

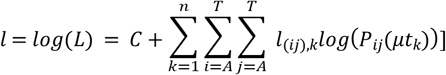

The maximum likelihood estimate cannot be obtained analytically. We relied on the Newton-Raphson method (26), which iteratively updates the new value of the mutation rate *u* until convergence.

The detailed derivations for both F81 and JC69 models can be found in the supplementary methods.

### Statistical analysis

Continuous variables were described using median and range; categorical variables were described using frequency and percentage. Demographics and variant characteristics were compared between patients in different virus groups by using ANOVA or Wilcoxon rank sum tests for continuous variables and Fisher’s exact or Pearson’s chi-square tests for categorical variables. The estimated mutation rates from two different groups are compared using the t-test, assuming the maximum likelihood estimates follow approximately a normal distribution. The confidence interval of the estimated mutation rate is calculated based on the maximum likelihood estimate following approximately a normal distribution N(u, 1/I(u)), where u is the true value, and I(u) is the Fisher information. PRISM software (version 8.4.3, GraphPad Software, San Diego, CA) and Python (version 3.7.4) with statsmodel package (version 0.13.2, for construction of ML models) was used for analysis.

## Results

From 3/17/2020 through 5/27/2020, a total of 40 paired nasopharyngeal samples (initial and subsequent) from acutely infected individuals with SARS-CoV-2 were identified and retrieved from the COVID19 biobank. Median days between paired tests was 13 days [IQR 8.5-20]. Median patient age was 54 years [IQR 31, 66] and included 20/40(50.0%) males with 26/40 (67.0%) being white, and with 28/40 (70.0%) having at least one comorbidity (table 1). Comorbidities included endocrine 23/40 (57.5%), cardiac 17/40 (42.5%), pulmonary 8/40 (20.0%) and Immune/Oncologic 6/40 (15.0%).

**Table 1.** Patient Demographics of Paired SARS-CoV-2 Isolates

|  | Total | wt NSP-14 | Δ NSP-14 | p-value |
| --- | --- | --- | --- | --- |
| Total pairs | 40 | 28 (70.0%) | 12 (30.0%) |  |
| Median interval (days) [IQR] | 13 [8.5, 20] | 13 [8.5, 20] | 14 [8.5, 20] | 0.72<sup>b</sup> |
| Demographics |  |  |  |  |
| Median Age (yr) [IQR] | 54 [31, 66] | 56 [31, 69] | 53 [32, 62] | 0.65<sup>b</sup> |
| Males | 20 (50.0%) | 14 (50.0%) | 6 (50.0%) | 0.99<sup>c</sup> |
| Race* |  |  |  | 0.46<sup>d</sup> |
| White | 26 (67.0%) | 16 (59.0%) | 10 (83.3%) |  |
| African American | 10 (26.0%) | 8 (30.0%) | 2 (16.7%) |  |
| Asian | 3 (7.5%) | 3 (11.0%) | 0 (0%) |  |
| Comorbidity |  |  |  |  |
| Any | 28 (70.0%) | 19 (67.9%) | 9 (75.0%) | 0.72<sup>d</sup> |
| Endocrine | 23 (57.5%) | 14 (50.0%) | 9 (75.0%) | 0.14<sup>c</sup> |
| Cardiac | 17 (42.5%) | 12 (42.9%) | 5 (41.7%) | 0.94<sup>c</sup> |
| Pulmonary | 8 (20.0%) | 5 (17.9%) | 3 (25.0%) | 0.68<sup>d</sup> |
| Immune/Oncologic | 6 (15.0%) | 4 (14.3%) | 2 (16.7%) | 0.99<sup>d</sup> |

\*Data not available for all subjects. Missing values: Race = 1.
Statistics presented as Median [P25, P75], N (column %).
P-values: b=Wilcoxon Rank Sum test, c=Pearson's chi-square test, d=Fisher's Exact test.

SARS-CoV-2 genomes of each pair were sequenced and mapped against the reference Wuhan strain (Wuhan-Hu-1, NC_045512.2). SNVs were identified for each pairing through IGV and filtered at allele frequencies (AF) ≥0.25, ≥0.5 and ≥0.75. A total of 120 SNVs changes between initial and subsequent samples were identified at AF ≥0.25, 53 at AF ≥0.5 and 33 at AF ≥0.75 (table 2). The majority of SNV changes were gained over the course of the infection (93/120 (77.5%), 32/53 (60.4%), 18/33 (54.8%) at AF ≥0.25, ≥0.5, ≥0.75 respectively) with the remainder being lost (27/120 (22.5%), 21/53 (39.6%), 15/33 (45.2%) at AF ≥0.25, ≥0.5, ≥0.75). Predominant SNVs were missense with most occurring in the ORF1a/b region and the spike protein region. While more SNVs were gained at low AF, there was no substantial difference between SNV types or gene location among different AF.

**Table 2.** Type and Location of SARS-CoV-2 Intra-host SNVs by Allele Fraction

| | AF ≥ 0.25 | AF ≥ 0.5 | AF ≥ 0.75 |
| --- | --- | --- | --- |
| **SNV changes** | 120 | 53 (44.2%) | 33 (35.0%) |
| **Mutations Gained** | 93 (77.5%) | 32 (60.4%) | 18 (54.8%) |
| **Mutations Lost** | 27 (22.5%) | 21 (39.6%) | 15 (45.2%) |
| **Missense** | 71 (59.2%) | 36 (67.9%) | 23 (69.7%) |
| **Silent** | 30 (25.0%) | 11 (20.8%) | 7 (21.2%) |
| **INDEL** | 2 (1.6%) | 2 (3.8%) | 1 (3.0%) |
| **Other** | 17 (14.2%) | 4(7.5%) | 2 (6.1%) |
| **ORF1 a/b** | 82 (68.3%) | 36 (67.9%) | 26 (61.9%) |
| **ORF3** | 4 (3.3%) | 3 (5.7%) | 3 (7.1%) |
| **ORF6** | 2 (1.7%) | 1 (1.9%) | 1 (2.4%) |
| **ORF7** | 1 (0.8%) | 0 (0%) | 0 (0%) |
| **ORF8** | 3 (2.5%) | 2 (3.8%) | 2 (4.8%) |
| **ORF10** | 1 (0.8%) | 0 (0%) | 0 (0%) |
| **Spike** | 16 (13.3%) | 6 (11.3%) | 5 (11.9%) |
| **Membrane** | 2 (1.7%) | 1 (1.9%) | 1 (2.4%) |
| **Envelope** | 0 (0%) | 0 (0%) | 0 (0%) |
| **Nucleocapsid** | 6 (5.0%) | 4 (7.5%) | 4 (9.5%) |
| **Untranslated region (UTR)** | 3 (2.5%) | 0 (0%) | 0 (0%) |

We identified 12/40 (30.0%) pairs with a non-synonymous mutation in NSP-14 (ΔNSP-14) while 28/40 patients (70.0%) did not (wtNSP-14). Median age, gender, race and comorbidities were similar between both groups. For both ΔNSP-14 and wtNSP-14 groups, the majority of SNVs were gained over the course of infection in both groups. Mutation types and locations were similar between groups (supplementary table 1 and 2).

Mutation rates were calculated through the F81 and JC69 models (figure 1, supplementary figure 1 for JC69). Focusing on F81 modeling, the mutation rate from all samples was found to be 93.6 substitutions/genome/year [95%CI 90.8-96.4] at AF ≥0.25, 40.7 [95% CI 38.9-42.6] at AF ≥0.5 and 34.7 [95%CI 33.0-36.4] at AF ≥0.75. Mutation rate of ΔNSP-14 were significantly higher at low AF compared to wtNSP-14 group (109.4 [95%CI 99.7-119.1] vs 86.0 [95%CI 82.1-89.9] substitutions/genome/year, p-value <0.001). Surprisingly, mutation rate was lower in ΔNSP-14 compared to wtNSP-14 both at AF ≥0.5 (32.0 [95% CI 26.8-37.2] vs 44.9 [95% CI 42.1-47.7] substitutions/genome/year, p-value <0.001) and at AF ≥0.75 (16.0 [95% CI 7.0-25.1] vs 39.8 [95% CI 25.0-54.5] substitutions/genome/year, p-value <0.001).

**Figure 1.**
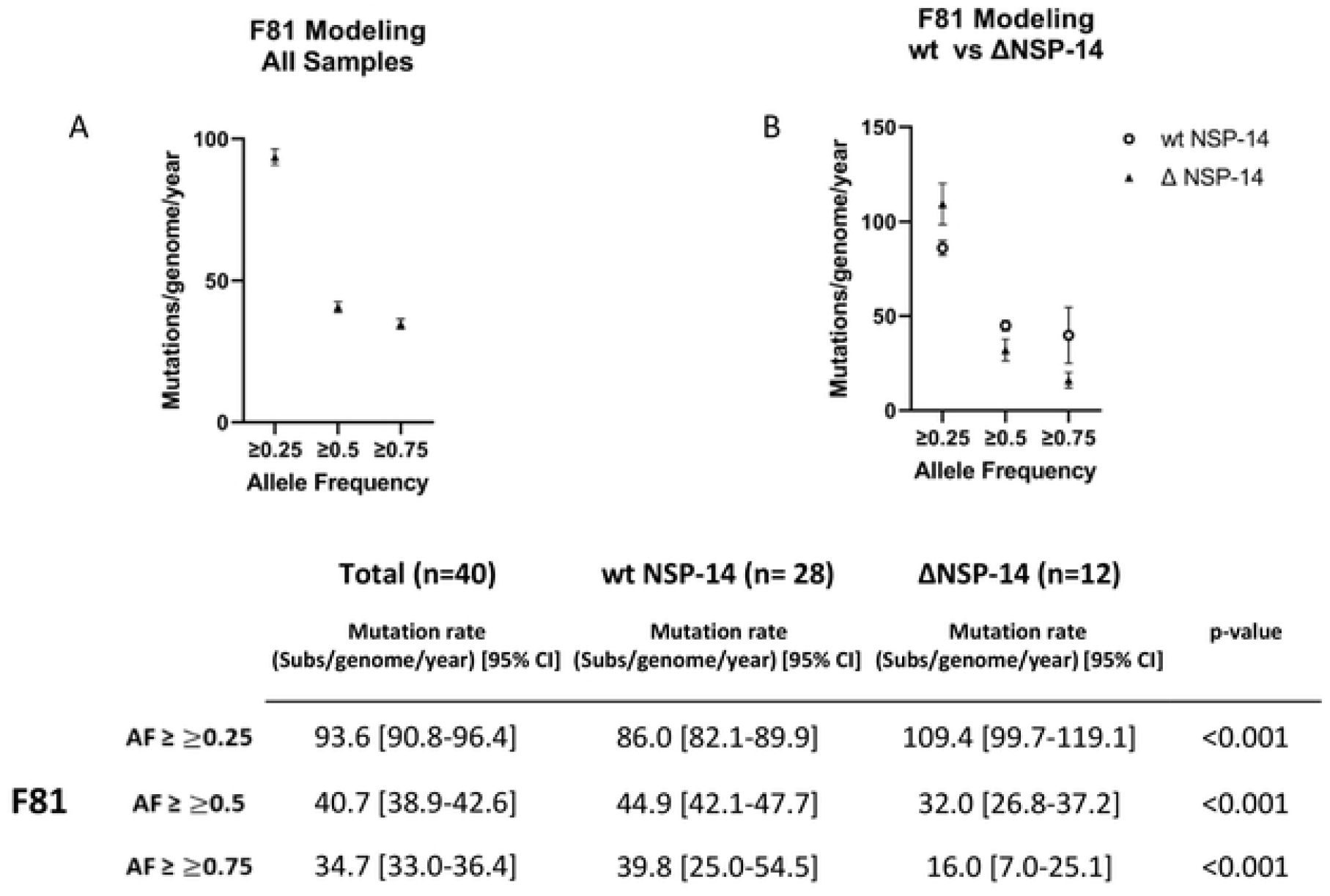
F81 Mutation Modeling by Allele Frequency with and without alteration in NSP-14. Graphic representation of F81 evolution modeling at AF ≥0.25, ≥0.5, ≥0.75 of A) total patient sample and B) comparison between wt and ΔNSP-14. Bars represent 95%CI. Table displaying data for F81 modeling is displayed below. P-values displayed represent comparison of wt and ΔNSP-14 groups.

Lastly, patients with underlying immunologic/oncologic comorbidities had a substantially higher mutation rate than other comorbidities at all three AF (figure 2, supplementary figure 2 for JC69). Mutation rate in patients with immunologic/oncologic comorbidities was 160 [95% CI 136.2-183.7] vs 81.2 [95% CI 78.1-84.2] substitutions/genome/year at AF ≥0.25, 137.9 [95% CI 115.8-160.0] vs 22.6 [95% CI 21.0-24.2] at AF ≥0.5 and 126.9[95% CI 105.7-148.0] vs 17.4 [95%CI 16.0-18.9] at AF ≥0.75. Overall mutation rates calculated through JC69 modeling were comparable to those with F81 at all three AF (supplementary figure 3). Results based on JC69 modeling are presented in Supplementary Figures 1 and 2.

**Figure 2.**
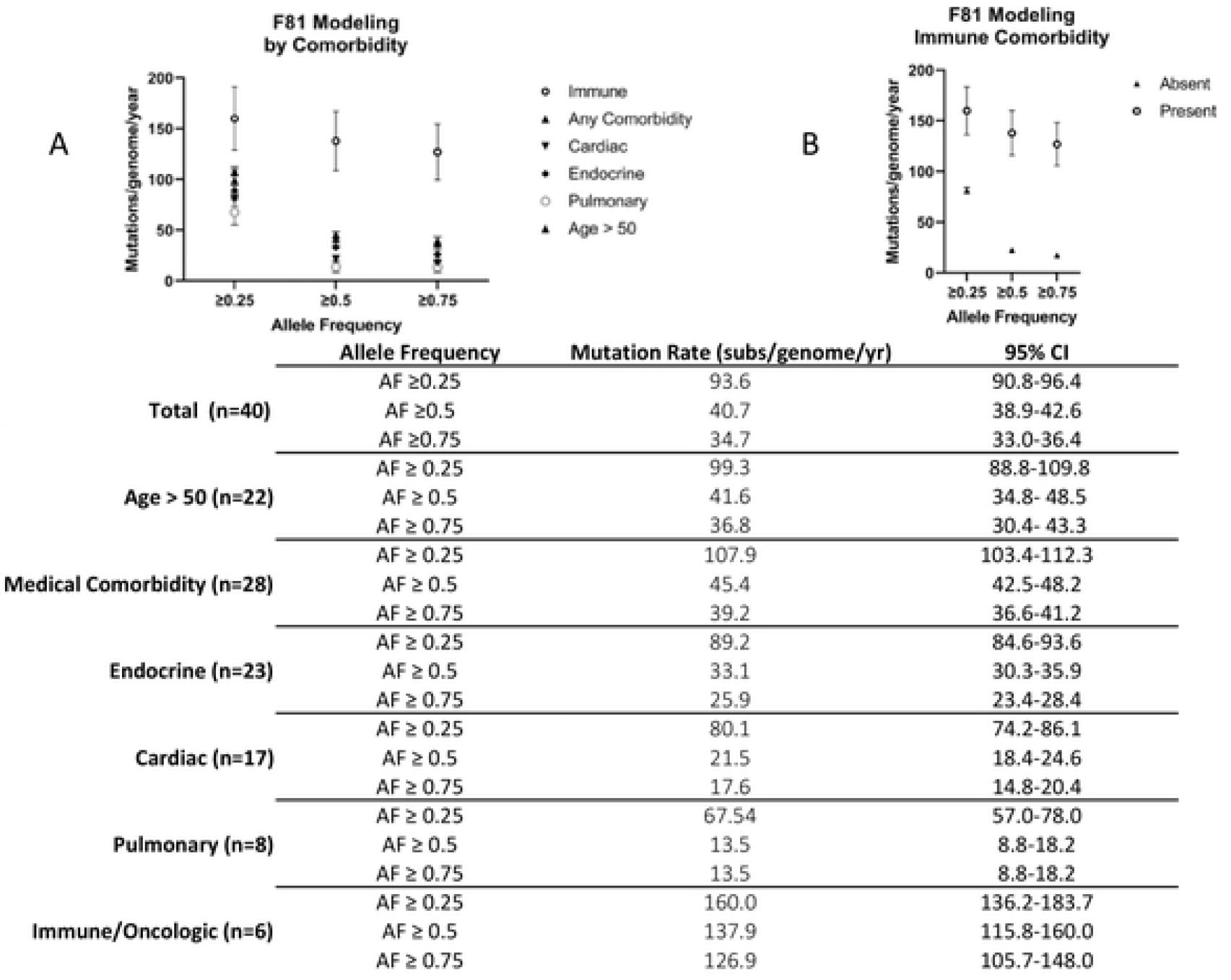
F81 Mutation Clock Modeling by Allele Frequency with Respect to Age and Comorbidity. Graphic representations of mutation rates at AF ≥0.25, ≥0.5, ≥0.75 for A) age and comorbidities and B) those with and without immunologic/oncologic comorbidity. Bars represent 95%CI. Table displaying data for F81 modeling is displayed below.

## Discussion

The dynamics of SARS-CoV-2 evolution remain poorly understood. The virus continues to change leading to the emergence of new variants adversely affecting pandemic response (27). The mutation rate commonly cited is calculated through analysis of unrelated regional and global sequences. These population based rates have ranged from 21.6 to 28.4 substitutions/genome/year (5). The rate of evolution of SARS-CoV-2 for much of 2020 was consistent with the virus acquiring approximately two mutations per month (28,29). However, recently the viral mutation rate has accelerated and now lies at its fastest point with the emergence of the Omicron variant (30).

Here, we analyze intra-host mutation rate at multiple allele frequencies to better characterize and understand the capacity for SARS-CoV-2 to evolve following its initial introduction and prior to external influence by antivirals, vaccinations and prior immunity. While intra-host mutation dynamics have been previously described (31), the intra-host mutation rate over the course of an infection, important for predicting future variant development has been poorly studied. We find the intra-host mutation rate is over 50% greater than what was reported through population based surveillance at AF ≥0.75 (the WHO standard). Additionally, if low frequency SNVs (<0.75) act as a reservoir for further generation of dominant mutations, the mutation rate can be up to 80% higher at AF ≥0.5 and nearly 350% greater at AF ≥0.25. Recognition of this mutation potential aids in our understanding of current evolutionary patterns and provides useful clues for future coronavirus pandemics (32,33).

By analyzing the genomic changes at lower AF, our study provides a better appreciation of intra-host SARS-CoV-2 biodiversity. We find the highest diversity at lowest AF (≥0.25) demonstrating that potential SNVs occur nearly 4 times higher than commonly reported. Fitness of these low frequency SNVs and their effect on transmission remains poorly understood. Current literature is skeptical of significant person to person spread of low AF SNVs and report only rare transmission recognized among individuals within the same household (6,7,34). However, it is reported that accelerated episodic increase in mutation rate (∼ 4 fold higher than the background substitution rate) drive the emergence of variants of concerns(35). We hypothesize that low AF SNVs may play a role in such a process.

Prior studies report that alteration in NSP-14 is associated with increased mutation load across the genome compared to other NSP changes (36). NSP-14 is vital for survival of various coronaviruses including SARS-CoV-2 (37). Inactivating NSP-14-ExoN in murine hepatitis virus (MHV-CoV) significantly altered recombination patterns and decreased recombination frequency compared with wild-type MHV-CoV (10). While virus diversity has been found to contribute to disease severity in coronaviruses including SARS-CoV-1 and MERS-CoV (32), further studies showed ExoN knockout mutants of MERS-CoV and SARS-CoV-2 are nonviable, suggesting excess mutation may have a deleterious effect (11,38). Our findings are consistent with this. While the mutation rate is significantly higher in ΔNSP-14, such change occurs only at low AF. This suggests SARS-CoV-2 viruses with altered NSP-14 may be less fit (37). As such, SARS-CoV-2 NSP-14 is being evaluated as a potential therapeutic target (10,12).

Lastly, SARS-CoV-2 genetic diversity and clinical outcome are influenced by host effects (33). High rates of mutation over short time periods have been seen in previous studies of immunosuppressed individuals chronically infected with SARS-CoV-2. (39–41). Additionally, prolonged viral shedding can occur in the immunocompromised population allowing for increased time to generate fit mutations (42). In one example, SARS-CoV-2 shedding was observed for as long as 471 days from the upper respiratory tract of a patient suffering from advanced lymphocytic leukemia and B-cell lymphoma. Throughout the course of this infection the accumulation of an unusually high number of immune escape mutations was detected and the mutation rate was calculated at 35.6 (95% CI: 31.6-39.5) substitutions per year through the Bayesian Skyline Model (43). In our study, we included patients with several comorbidities, only viruses originating from hosts with immune comorbidities were found to have significantly accelerated mutation rate (44). This adds to the growing understanding that a patient’s immunity profile impacts viral evolution over the course of the infection (43). Better delineation of specific immune factors associated with alteration of evolutionary rate are needed.

There are several limitations to this study. First, while our investigation of 40 SARS-CoV-2 patient pairs demonstrated substantially higher mutation rate than commonly reported, further analysis with larger cohorts would improve accuracy. Similarly, patients were grouped in broad comorbidity categories rather than by more specific underlying disease. Studies with greater characterization of underlying comorbidities, particularly immune, will provide a better picture of host factors associated with alteration in SARS-CoV-2 mutation (42,45). While a cutoff AF ≥ 0.75 was based on WHO guide for global variant surveillance, the significance of lower frequency SNVs remains unclear. This study sheds more light on the virus diversity identified at lower AF thresholds. By focusing analysis on viral isolates originating from the initial pandemic wave, ours is the first study to determine the intra-host mutation rate of SARS-CoV-2 prior to the influence of many external factors (e.g. antiviral medications, monoclonal antibody therapy, immunization, and natural immunity from prior infection). Determining the effect of pharmacologic interventions, immunization and previous infection on the mutation rate of subsequent SARS-CoV-2 isolates is a logical next step. Additionally, analysis of subsequent SARS-CoV-2 variants (Alpha, Delta, and Omicron) with parameter rich models such as HKY or GTR are currently being planned. Lastly, placement of patients within wt and ΔNSP-14 groups occurred without association to gene location or type. It is possible that several NS mutations placed in this group did not substantially affect NSP-14 function. Further study focusing on those SNVs with a defined effect on NSP-14 activity are needed (45).

## Conclusion

Our study demonstrates the intra-host mutation rate of SARS-CoV-2 is substantially higher than previously reported through population based analysis. In addition, low frequency intra-host mutations may be an important reservoir contributing to possible future variant emergence. SNVs in NSP-14 were found to have increased mutation rate but only at low AF. Conversely, we find enhanced mutation rate in immunocompromised patients while no elevation was observed in patients with underlying cardiac, pulmonary or endocrine comorbidities. SARS-CoV-2 intra-host dynamics have crucial implications on current and future pandemic planning, development of vaccines, and antiviral therapy.

## Authors contributions

K E H, FE, and BR conceptualized and directed this research. TA, XL, XZ and JL, developed methodology, and performed evolutionary modeling and mutation statistics. TJ and YC assisted in sample acquisition, Illumina sequencing and pipeline development. DR and JK assisted in study design, sample identification and acquisition. SW assisted in statistics review. All authors contributed to discussions and manuscript preparation.

## Acknowledgments

We appreciate Daniel H. Farkas, PhD, for his kind insight and thoughtful review of the project.

